# The wildlife-livestock interface modulates anthrax suitability in India

**DOI:** 10.1101/419465

**Authors:** Michael G. Walsh, Siobhan M. Mor, Shah Hossain

**Affiliations:** The University of Sydney, Faculty of Medicine and Health, Marie Bashir Institute for Infectious Diseases and Biosecurity, Westmead, New South Wales, Australia; The University of Sydney, Faculty of Medicine and Health, Westmead Institute for Medical Research, Westmead, New South Wales, Australia; University of Liverpool, Faculty of Health and Life Sciences, Institute of Infection and Global Health Liverpool, Merseyside, United Kingdom; The University of Sydney, Faculty of Science, School of Veterinary Science, Camperdown, New South Wales, Australia; Prasanna School of Public Health, Manipal Academy of Higher Education, Manipal, Karnataka, India

**Keywords:** anthrax, landscape epidemiology, infection ecology, wildlife-livestock interface, India

## Abstract

Anthrax is a potentially life-threatening bacterial disease that can circulate in wild and domestic animals and subsequently spillover to human contacts with devastating consequences for human and animal health, as well as livestock economies and ecosystem conservation. India has a high annual occurrence of anthrax in some regions, but a country-wide delineation of risk has not yet been undertaken. The current study modeled the geographic suitability of anthrax across India and its associated environmental features using a biogeographical application of machine learning. Both biotic and abiotic features contributed to risk across multiple scales of influence and the wildlife-livestock interface, using elephants as a wildlife sentinel species, was the dominant feature in delineating anthrax suitability. In addition, water-soil balance, soil chemistry, and historical forest loss were also influential. These findings suggest that the wildlife-livestock interface plays an important role in the cycling of anthrax in India. Prevention efforts targeted toward this interface, particularly within anthropogenic ecotones, may yield successes in reducing ongoing transmission between animal hosts and subsequent zoonotic transmission to humans.

## Introduction

Anthrax is a global disease of tremendous impact to the economy and health of pastoralist communities. It is responsible for substantial mortality in livestock in temperate and tropical settings, and can result in spillover to humans in contact with livestock and wildlife resulting in life-threatening cutaneous, gastrointestinal or respiratory disease(1). Risk of human infection is greater in those that process or consume contaminated animal products, bushmeat and carcass meat. However, humans are accidental dead-end hosts and generally do not contribute to onward transmission of the disease. Sporadic, epizootic transmission occurs in wildlife also and can have devastating impacts, especially among mammalian herbivores(2). The causative agent, *Bacillus anthracis*, can remain inactive, yet stable, in soil for up to several decades and across a spectrum of environmental conditions owing to the generation of spores, which form upon contact with atmospheric oxygen. In a favourable environment these spores germinate into the vegetative form and replicate. For example, following ingestion by grazing mammals, spores germinate and begin replicating in the newly infected host(1,3,4). External hemorrhage from multiple orifices is a common clinical feature in animals at the time of death and leads to continuation of the cycle through re-contamination of the environment with bacteria and subsequent grazing of susceptible herbivores therein.

Anthropogenic pressure operates within several spheres of influence in domestic and sylvan landscapes to promote disease emergence(5). Two such spheres of influence, the wildlife-livestock interface and habitat loss, may be particularly important in propagating the anthrax infection cycle. The wildlife-livestock interface, defined as the extent to which wildlife species and livestock interact (or have the potential to interact) directly or indirectly in a given landscape, has been recognized for some time as driving inter-species transmission between animals (Figure 1), and subsequent potential spillover to humans(5–12). The wildlife-livestock interface has been shown to significantly enhance the ability of anthrax to persist in some areas, and may be a key driver of enzootic transmission in tropical settings(6,8,13). Habitat loss, particularly over long periods of time, can substantially alter the abundance, richness, and movement patterns of the wildlife species that occupy the transitional spaces, or ecotones(5,13–18). This in turn can directly or indirectly influence the wildlife-livestock interface(10,12). Once a wildlife-livestock interface has been established in an ecotone, the characteristics of *B. anthracis* are such that inter-specific transmission can modulate plant-animal interaction. For example it has been shown that anthrax-infected carcasses promote grasses favored by grazers, which subsequently draw these species to the spore-containing soils(19,20).

**Figure 1.**
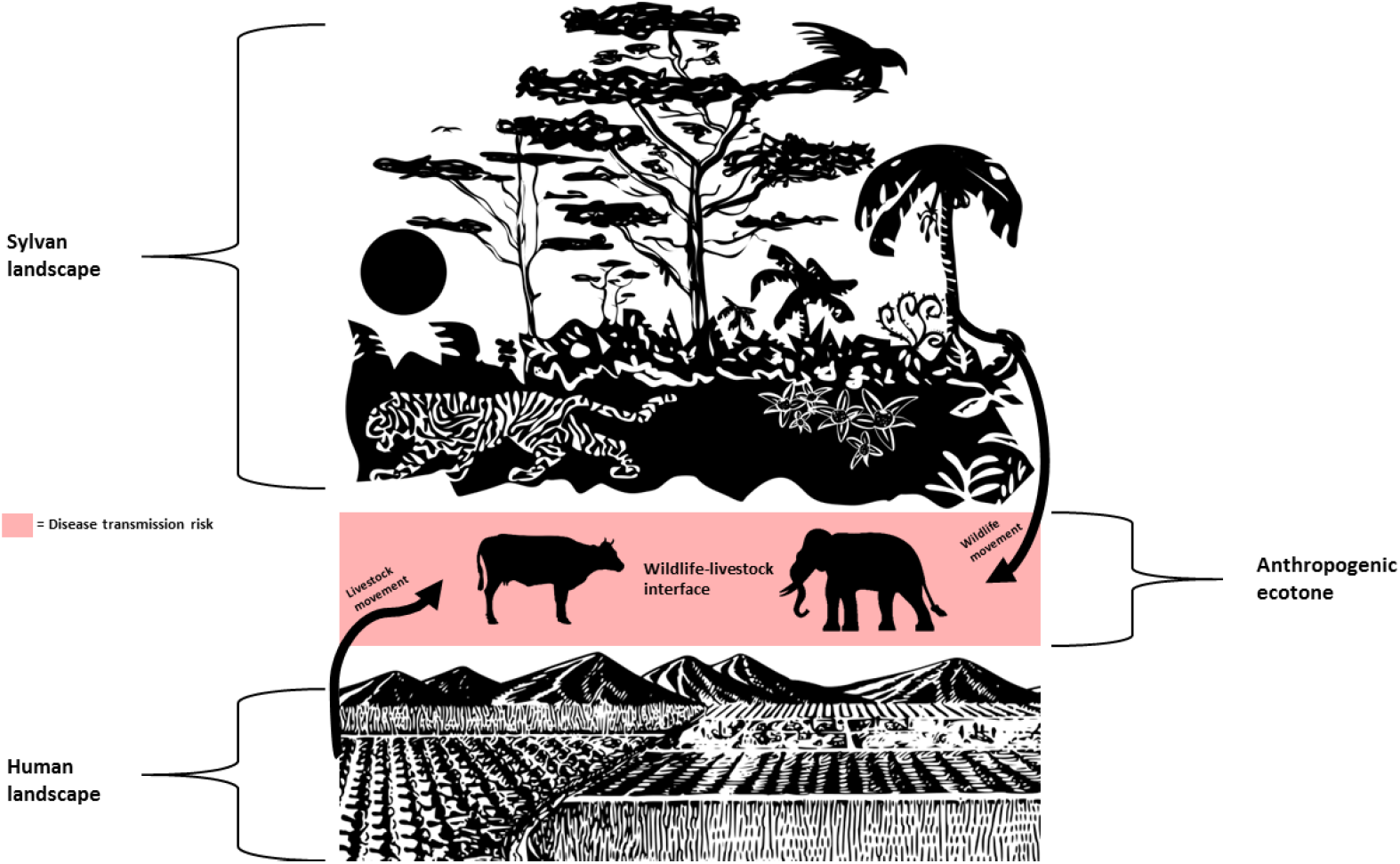
Representation of the wildlife-livestock interface in anthropogenic ecotones.

India has a high burden of anthrax, with some regions experiencing annual or near annual outbreaks of disease(21,22). A significant number of these outbreaks occur in wildlife, with a preponderance of reported cases in elephants(7,23–26). Common points of interface in Indian settings include shared waterholes, salt licks and grazing meadows, particularly in the forest fringe areas. These also tend to be the areas where backyard animal husbandry, comprised of small and mixed holdings of grazing animals, are in closest proximity to wildlife. The over-representation of elephants may be due to their unique biological and ecological requirements in landscapes increasingly shared with humans and their livestock, or may simply reflect their role as a sentinel species. Typically in the wild, dying or dead animals are quickly devoured by predators or scavengers. Thus detection of anthrax cases in wildlife is often difficult, because they are cleared before samples can be obtained. However, elephants are a notable exception because of their size. Anthrax surveillance is not consistent across the country and the data generated is not always reliable. The current investigation sought to map geographic anthrax suitability as a representation of epidemiological risk, while inferring ecological relationships between anthrax outbreaks and abiotic and biotic features. Specifically, we hypothesized that an increasing wildlife-livestock interface, as marked by a key sentinel wildlife species (elephants), and proximity to human-modified landscapes (specifically, historical forest loss) would delineate greater anthrax suitability.

## Methods

One hundred and three anthrax outbreaks between 1 January, 2000 and 1 May, 2018 were identified from the ProMED-mail electronic surveillance system. This system is maintained by the International Society of Infectious Diseases and provides archival documentation of formal and informal reports of infectious disease occurrences[25]. ProMED-mail reports are screened by a large team of editors, moderators, correspondents and sometimes country managers, who evaluate reports as well as engage the large group of locally-sourced subscribers to offer additional insight in support of or against the alerts(27). The data thus represent a specific cross-section of disease experience rather than a population-based sample. As such, while we emphasize that the scope of this study does not apply to the full spectrum of anthrax experience in India, we do correct for potential reporting bias inherent in the data (see statistical methods below), which minimizes reporting bias in the assessment of anthrax suitability in anthropogenic environments. In addition, we also evaluated model performance using an independent sample of 22 laboratory confirmed anthrax outbreaks, with investigations reported in the scientific literature (n = 17) (23,28–38) or by government agencies (n = 5) as captured by the EMPRES Global Animal Disease Information System (EMPRES-i; http://empres-i.fao.org), which is maintained by the Food and Agriculture Organization. This latter evaluation (see statistical methods below) has the added benefit of providing the first validation testing of ProMED-mail surveillance data in India to delineate risk of an important zoonotic infection.

The Gridded Livestock of the World (GLW) provided livestock densities for cattle, sheep, and goats at 30 arc second resolution (approximately 1km) (39). The data thus obtained are more up to date than the periodic animal census data available for India. Because the current aim was to assess anthrax suitability associated with livestock presence, a combined livestock raster was created based on the sum of the absolute number of cattle, sheep, and goats per unit area, rather than calculating livestock units(40) because evaluating the differential impact of different livestock species on the grazing and browsing capacity of land parcels was beyond the scope and capacity of this study given the lack of sufficiently fine-scaled environmental or outbreak data. The Global Biodiversity Information Facility (GBIF) was used to identify observed free-ranging elephants (*Elephas maximus*) across India so their ecological niche could be modeled and used as a wildlife sentinel species (http://www.gbif.org/). These two data products, livestock density and the elephant niche, were then applied to the quantification of the wildlife-livestock interface (see statistical methods description below). The elephant niche was chosen as the representative wildlife niche because of the species’ importance as both an anthrax sentinel(7,23–25,41) and the common overlap of their range with that of grazing livestock in forest fringe areas(29). However, the utility of this niche could not be compared to that of other species because of a corresponding lack of adequate observations of other species occurrences in GBIF. Climate data were obtained from the WorldClim Global Climate database(42). The mean annual temperature was calculated using aggregate spatio-temporal weather station data between 1950 and 2000, and extracted as a 30 arc second resolution raster(43). The Priestley-Taylor α coefficient (P-Tα) was used to represent water-soil balance(44,45). The P-Tα is the ratio of actual evapotranspiration to potential evapotranspiration, and captures water availability in the soil, as well as the local vegetation’s water requirements, as contrasted with solar energy input. Thus, P-Tα is a robust estimate of environmental water stress through soil-water balance. The raster data was acquired at 30 arc seconds resolution from the Consultative Group for International Agricultural Research (CGIAR) Consortium for Spatial Information. The ratio is dimensionless and ranges from 0 (extreme water stress) to 1 (no water stress)(46).

Soil pH and organic content data were obtained from the Global Soil Dataset for Earth System Modeling, which is based on an improved protocol of the Harmonized World Soil Database(47). These two rasters are recorded at 5 arc minutes (approximately 10 km).

Historical forest data from 1900 were derived from the History Database of Global Environment (HYDE) anthrome data product at 5 arc minutes resolution and compared to forest cover in 2000(48,49). The HYDE database is the result of an ongoing development by the Netherlands Environmental Assessment Agency to describe human population growth and land use change from 10,000 BCE to the present day. Estimation of land cover and land use is based on an amalgam of satellite image data, historical sub-national statistical inventories, ecosystem envelopes based on climate and geological (soil) properties, and archeological data(48,49). The difference in proximity to historical forest cover (i.e. forest cover that was present in 1900) and modern forest cover (i.e. forest cover present in 2000) was thus calculated and evaluated with respect to anthrax suitability.

The sampling of background points was weighted using the human footprint (HFP) to correct for potential reporting bias in anthrax presence points (see modeling description below). The HFP was quantified using data obtained from SEDAC(50), and is comprised of two stages(51). First, the human influence index (HII) describes the impact of human influence based on eight domains: 1) population density, 2) proximity to railroads, 3) proximity to roads, 4) proximity to navigable rivers, 5) proximity to coastlines, 6) intensity of nighttime artificial light, 7) location in or outside urban space, and 8) land cover. These domains are scored and quantify the level of human impact per item per 1km^2^. The eight domains are then combined to form a composite index, which ranges from 0, indicating an absence of human impact, to 64, indicating maximal human impact. The HII is then normalized to the 15 terrestrial biomes defined by the World Wildlife Fund to obtain the HFP. The normalization is the ratio of the range of minimum and maximum HII in each biome to the range of minimum and maximum HII across all biomes, and is expressed as a percentage(51).

## Statistical Analysis

Anthrax suitability in India, as well as the ecologic niche of elephants, was modeled using maximum entropy (Maxent) machine learning. Machine learning is now widely applied to the modeling of the geographic suitability of many zoonoses(52–54), and Maxent is an analytically appealing approach because the model is not constrained by a specific functional form. Additionally, the system can be modeled without knowledge of the locations of unknown (and unknowable) anthrax outbreak absences(54,55). Maxent has become a popular implementation to ecological niche modeling due to its robustness(56).

A metric for the wildlife-livestock interface was constructed by weighting the density of all livestock in each 1 km^2^ space by the ecological niche of *E. maximus*, which was estimated by a separate Maxent model using observations of *E. maximus* from the GBIF. The new weighted metric raster was the product of the livestock density raster and the probability density function raster of the estimated niche. As such, the interface is a representation of the number of livestock present in a given space adjusted by the probability of that space being suitable habitat for elephants.

Annual temperature, P-Tα, soil chemistry, the difference in forest cover between 1900 and 2000, the wildlife-livestock interface, and livestock density were included in the Maxent model of anthrax suitability. All covariates were aggregated to scales of 5 and 30 arc minutes, respectively, for the two spatial scales modeled separately in this study (see below). Correlation was low (all values of ρ were < 0.6) among the included environmental covariates. A total of 10,000 background points were sampled, weighted by HFP as described above to correct for reporting bias in anthrax surveillance(57). Five-fold cross-validation was used to train the model, and the area under the receiver operating characteristic curve (AUC) was used to evaluate model performance. To correct for overfitting, the regularization parameter was set at 1.0. Covariates were ranked by permutation importance, which is a random permutation of their values between background and presence points during training(54,58). With all available covariates under consideration, the best model was selected based on performance and fit using 1) the test AUCs between the full and reduced models, 2) the Akaike information criterion (AIC) based on a Poisson point process to measure goodness-of-fit(59), and 3) a jackknife variable selection procedure wherein each covariate’s single contribution to the loss function is compared to the loss function when the covariate is withheld from the model. Because the surveillance data were derived from the ProMED system, the model performance was evaluated using the independent testing dataset described above, which is comprised of a separate independent sample of laboratory-confirmed anthrax outbreaks. Reported AUCs, therefore, reflect the externally validated model performance.

As some recent work identified, the relationship between infectious diseases and abiotic and biotic features may depend on spatial scale(60). We therefore include two sensitivity analyses to evaluate whether these features operate differently at different scales. First, the models were evaluated at 5 arc minutes and 30 arc minutes to determine whether predicted anthrax suitability was robust to scale and whether the abiotic and biotic features in the model demonstrated scale-dependence. Second, two additional reduced models, one abiotic (climate, soil chemistry, and forest loss) and one biotic (livestock density and the wildlife-livestock interface), were compared to the best fitting model with respect to model performance (AUC), model fit (AIC), and niche equivalency. The latter is an assessment of the degree to which predicted niches are coincident with each other(61). The maxent function (dismo package; v. 0.9-3) was used to fit the models(55,62,63). All analyses were performed using R statistical software version 3.1.3(64). The vector images used in Figure 1 are public domain content obtained from https://publicdomainvectors.org/en/ and distributed without restriction under Creative Commons CC0 1.0 Universal.

## Results

The distribution of reported ProMED-mail (red), and independent laboratory-confirmed (black), anthrax outbreaks in India from 1 January, 2000 to 1 May, 2018 is presented in Figure 2, while the distributions of environmental features are presented in S1 Figure 1.

**Figure 2.**
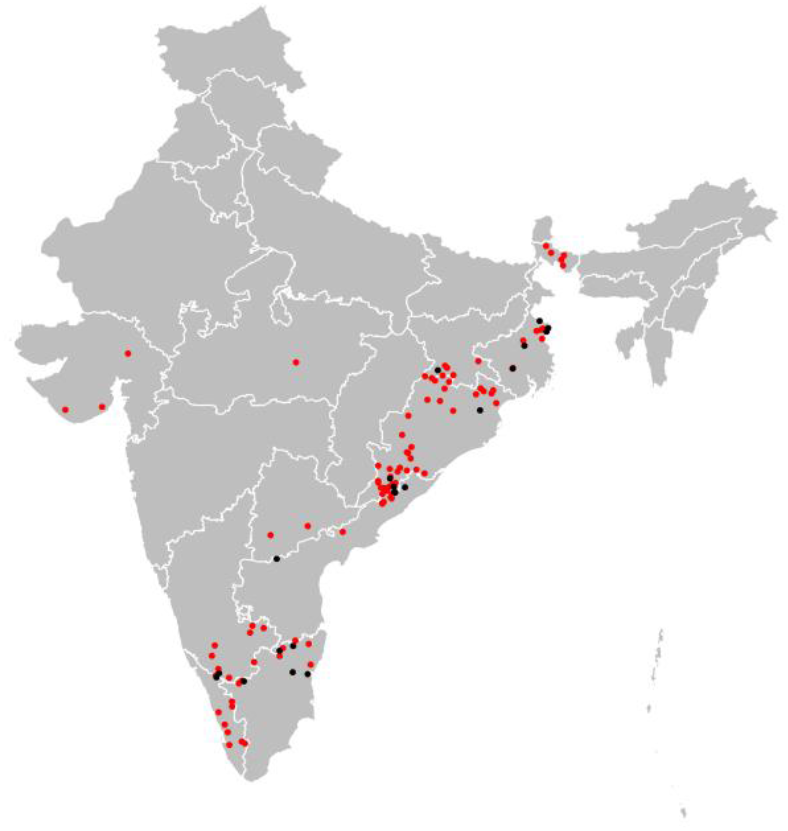
Distribution of anthrax outbreaks documented by the Pro-MED mail surveillance mechanism between 2000 and 2018 (red) and an independent sample of laboratory-confirmed outbreaks (black) used for model testing and evaluation. All maps are displayed only for the purposes of depicting the distribution of disease occurrence and risk, and do not reflect the authors’ assertion of territory or borders of any sovereign country including India. All maps created in R (v. 3.3.1).

The best fitting and performing model from all those considered (Model 6, S2 Table 1; S3 Figure 2) is presented in Figure 3. The wildlife-livestock interface was the dominant feature delineating anthrax suitability (permutation importance (PI) = 67.7%). Livestock density (PI = 10.5%), soil-water balance (PI = 7.1%), soil pH (4.7%), and proximity to forest lost between 1900 and 2000 (PI = 4.0%) were also influential to anthrax suitability. When the model was validated against the independent testing data, it performed reasonably well with AUC equal to 88%, and demonstrated the best fit (lowest AIC), compared to the other models (S2 Table 1). The jackknife variable selection model largely agreed with the final model above with respect to variable importance (S4 Figure 3), although livestock density appeared to contribute the least information to the model by itself and to have relatively little information present that is not already present in other variables. Therefore, a further reduced model with livestock density omitted was also considered (Model 7, S2 Table 1). The jackknife results notwithstanding, the reduced model with livestock omitted performed only slightly worse, but provided a noticeably poorer fit and so the model with livestock density included was retained as the final model. Given the strong association with the wildlife-livestock interface, which used the ecological niche of *E. maximus* as the wildlife sentinel with which to construct the interface, there was concern that the model fit and performance may be driven by elephant anthrax outbreaks specifically. As a sensitivity analysis, the twenty elephant outbreaks were removed and the Maxent model refit to the remaining 83 outbreaks. This model was similar in performance (AUC = 87%) and fit (AIC = 224) and demonstrated exceptional overlap in suitability (niche equivalency = 0.99; S5 Figure 4), suggesting that the inclusion of elephant outbreaks was not responsible for the close association between anthrax suitability and the wildlife-livestock interface.

**Figure 3.**
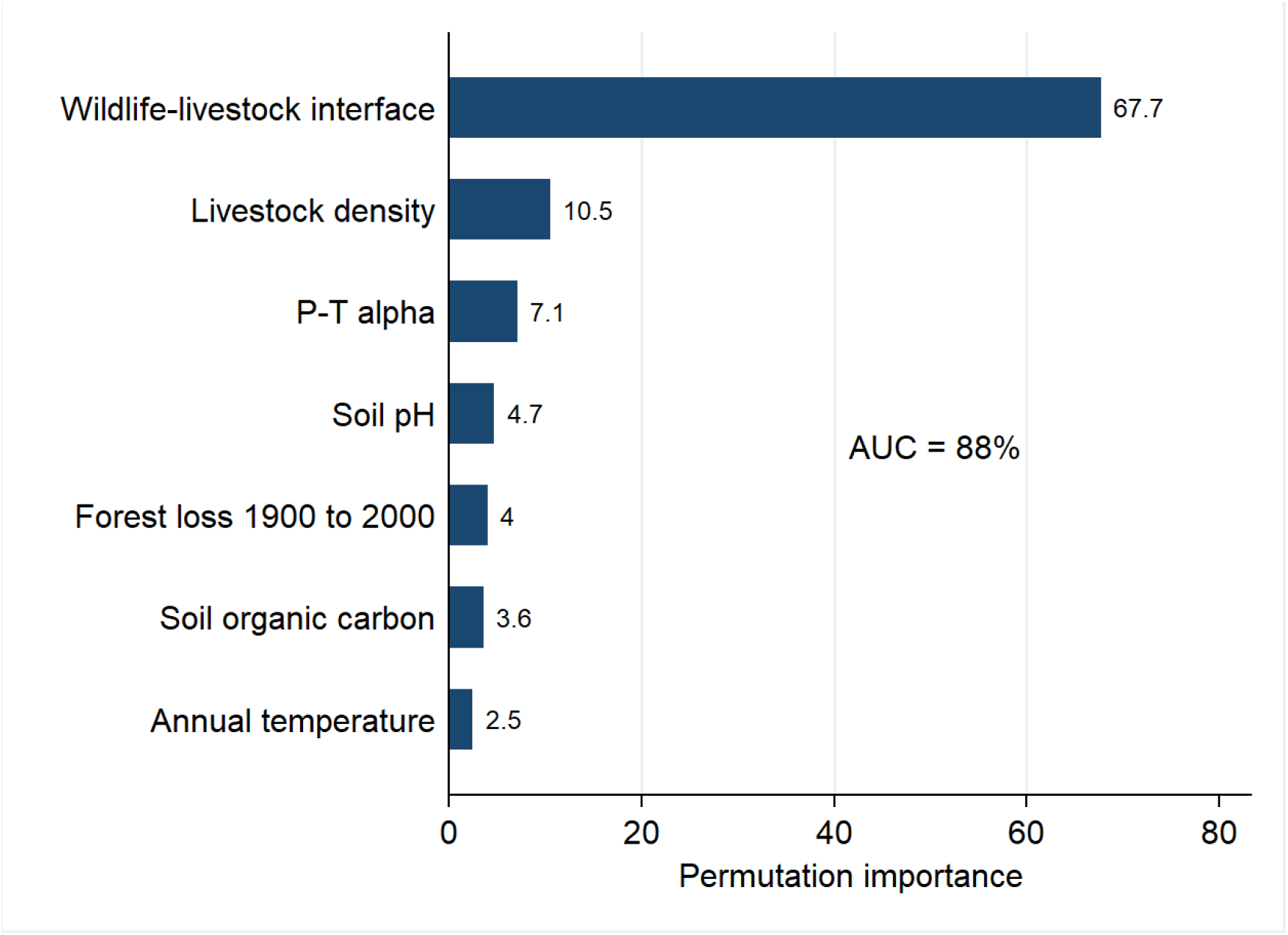
Environmental feature ranking by permutation importance in the Maxent model. The area under the curve (AUC) is reported as a percentage.

**Figure 4.**
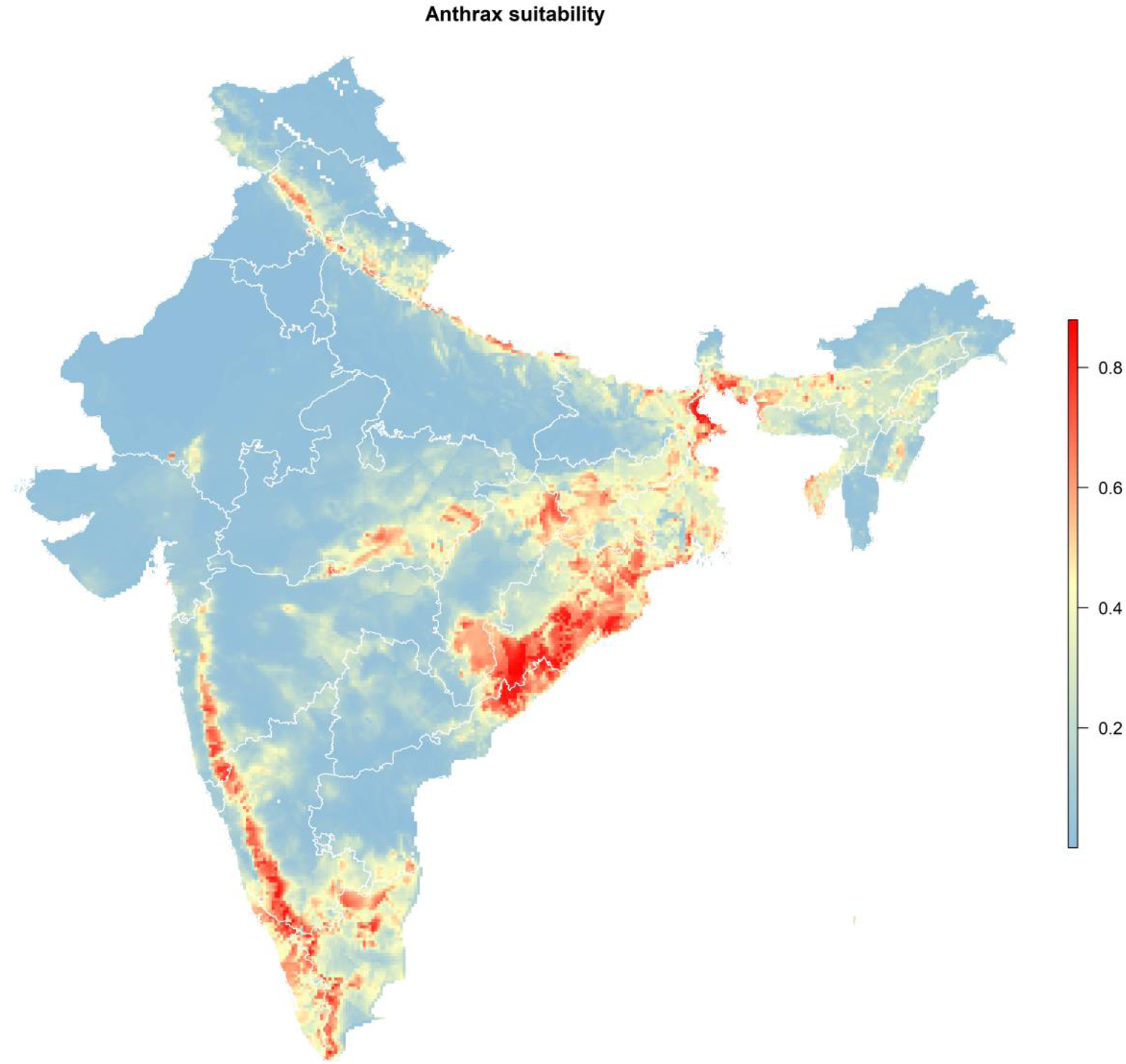
Predicted geographic anthrax suitability across India based on the best fitting and performing Maxent model (Model 6). All maps are displayed only for the purposes of depicting the distribution of disease occurrence and risk, and do not reflect the authors’ assertion of territory or borders of any sovereign country including India. All maps created in R (v. 3.3.1).

Finally, the wildlife-livestock interface did not appear to operate differently with respect to anthrax suitability at different scales (S2 Table 1, Model 6: 5 arc minutes vs. 30 arc minutes), while soil pH and forest loss did appear to exhibit a moderately increased impact at smaller scale (30 arc minutes) than at larger scale (5 arc minutes). This scale dependence was further confirmed by the abiotic model (S2 Table 1, Model 6: 5 arc minutes vs. 30 arc minutes), which evaluated these features independently of livestock density and the wildlife-livestock interface. Despite the moderate scale dependence observed for some of the abiotic features, the predicted suitability equivalency remained high when comparing the full model to the biotic model (niche equivalency at 5 arc minutes = 0.97; niche equivalency at 30 arc minutes = 0.97) or comparing the full model to the abiotic model (niche equivalency at 5 arc minutes = 0.95; niche equivalency at 30 arc minutes = 0.95).

Predicted anthrax suitability across India based on the best fitting and performing model is displayed in Figure 4. Two distinct corridors emerged: a wide distribution of high suitability running from northeast Andhra Pradesh to West Bengal in the east, and a narrower distribution of high suitability running the length of the Western Ghats and their ecotonal fringes from Kerala and Tamil Nadu north to southern Maharashtra in the west.

Figure 5 presents the response curves depicting the functional relationship between anthrax suitability and each environmental feature conditional on all others. Anthrax suitability was highest with soil pH in the range of 6 – 8 and soil organic carbon content in the range of 1% - 2%. Suitability increased with increasing soil-water balance until peaking at P-Tα = 0.6, which signifies low water stress. Anthrax suitability increased sharply with an increasing wildlife-livestock interface. Finally, increasing proximity to lost forest was associated with increasing anthrax suitability, with peak suitability within a window of 10 km of forest loss between 1900 and 2000.

**Figure 5.**
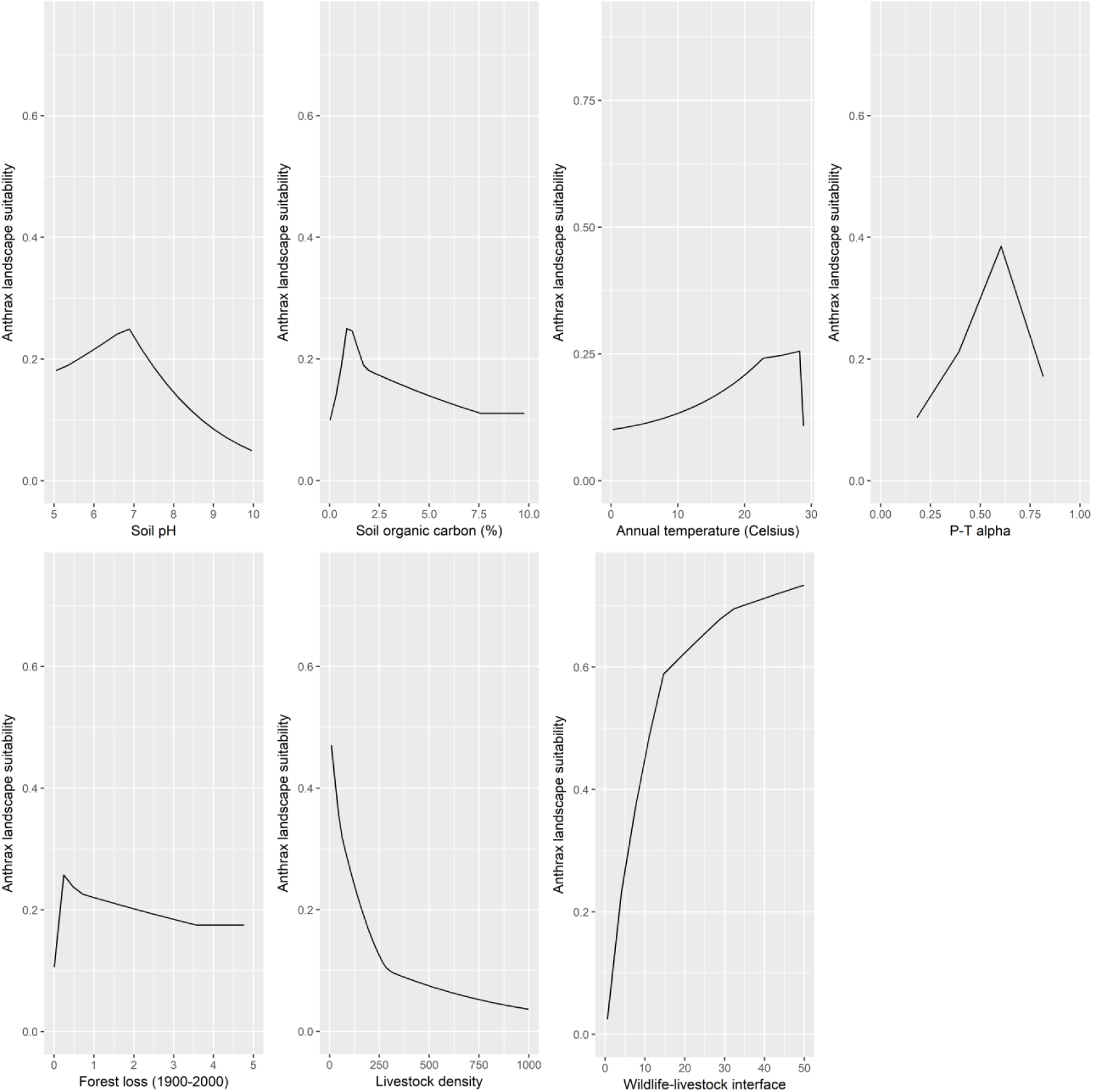
Response curves showing the functional relationships between each feature and anthrax suitability.

## Discussion

This study presents the first description of anthrax suitability across India. The wildlife-livestock interface was the most influential feature to anthrax suitability at relatively large and small scales, and even after removing wildlife cases (elephants) from the model. Moreover, this finding was consistent with the highest anthrax suitability manifesting in relatively small livestock herds, which would be more likely to occupy forest fringe than large industry herds. Soil-water balance and soil chemistry were also impactful to anthrax suitability, as expected. In addition, location within an approximate 10 km window of historical forest that has since been lost was moderately associated with increased suitability.

The relationship with the wildlife-livestock interface is unsurprising since it has been shown that intense competition for resources exists between livestock and wild grazing and browsing species, especially elephants, in precisely the same areas as those identified as high risk in the current study(29). Moreover, relatively small free ranging livestock herds will present the greatest opportunity for interface with sylvan species in anthropogenic ecotones. This was evident in the current study wherein greater anthrax suitability was associated with smaller herds. Both the Eastern and Western Ghats have stretches of forest covering several states with free ranging wild elephants through known corridors. These forests are often occupied by human habitations, particularly traditional forest-living people, for whom this wildlife-livestock interface is a constant reality. Similar wildlife-livestock interface foci have been identified in anthrax hotspots in other tropical settings as well(6,8). In the current study *E. maximus* was used as the sentinel wildlife species with which to adjust livestock density and thus quantify the wildlife-livestock interface. As such, this interface represents a sentinel interface rather than a complete interface, and therefore precludes the evaluation of a more varied interface between livestock and wildlife species. However, we feel this is an appropriate sentinel given the overwhelming preponderance of elephants among the wildlife outbreaks (>95%). Moreover, *E. maximus* was the only species for which sufficient data were available in GBIF to estimate a wildlife niche.

Soil pH in this study was generally reflective of the preferred pedological profile of *B. anthracis* with suitability peaking in the 6-8 range, which is typical for these bacteria in many parts of the world(3,4,65,66). In contrast to temperate climate regimes(67), anthrax suitability peaked at the lowest levels of organic carbon content, but this was likely indicative of the relative homogeneity of generally low soil organic carbon across much of India. Similar homogeneity was observed for annual temperature but not soil-water balance. The relationship with soil-water balance was interesting because, while the current study identified decreased water stress to be associated with greater anthrax suitability in India, increased water stress was associated with greater suitability across the temperate zones of the northern hemisphere(67).

The association between anthrax suitability and forest loss, albeit considerably weaker than the wildlife-xlivestock interface, is also intuitive and likely an important modulator of the interface. This novel finding may suggest a “revenant” forest presence that reflects historical cycling of anthrax between wildlife and livestock, or it may simply reflect a more generic increased proclivity to modern cycling in transformed sylvan landscapes. Large mammalian herbivores require extensive forest range in which to graze or browse. Elephant caloric requirements, for example, represent an extreme in plant intake and consequent home range. Adults will consume an average of 150 kg of vegetation per day, and can forage a variety of plants comprising 75 species across a range of 25 km^2^ (monsoon season) to 64 km^2^ (dry season) in southern India(68). The loss of forested habitat across India has reduced the available sylvan range for elephants, forcing them increasingly into anthropogenic ecotones wherein the potential for inter-specific contact is substantial(69,70). It is, therefore, expected that these transitional zones will reflect a trajectory of forest loss and, given the ability of spores to persist for decades in the environment, potential for long term cycling of anthrax between wildlife and livestock. However, the current data are too limited in temporal granularity to make this claim definitely, as anthrax cycling could also reflect a more recent introduction into ecotonal areas by relatively new livestock herds. Nevertheless, the association with the wildlife-livestock interface combined with close proximity to historical forest loss suggests the protection of wildlife populations and forest management, with concerted effort to maintain separation between wildlife and livestock, may be a fruitful approach to anthrax prevention in those areas of highest risk in India.

This study has some limitations that warrant further discussion. First, the anthrax outbreaks used to train the models in this study are based on ProMED-mail surveillance, which we recognize does not identify all outbreaks. In particular, the responsiveness potential of the surveillance system varies by state according to the quality of veterinary services and reporting infrastructure. This may lead to reporting bias in the identified anthrax locations. We attempted to correct for such reporting bias by selecting background points weighted by the presence of the human footprint as a proxy for reporting infrastructure. In addition, we tested the fitted model against an independent laboratory-confirmed sample of anthrax outbreaks to provide a less biased assessment of performance. Nevertheless, we concede that the data may not be representative of the complete anthrax experience in India over the last two decades. Second, even at the larger of the two scales considered here, the scale of the study is coarse by virtue of the scale of ProMED-mail reporting and the available environmental data. While this is unlikely to be of substantial influence to abiotic environmental features, which are expected to dominate at small (i.e. coarse) spatial scale, it may be influential to biotic features, which are expected to dominate at large (i.e. fine) scale(60). Third, climate features (PT-α and mean annual temperature) were constructs of decadal averages from 1950 to 2000, and therefore the models presented here assume temporal homogeneity of these aggregates both during the period in which they were recorded as well as during the period of observed anthrax outbreaks under current investigation.

In conclusion, this study has provided the first country-wide predictions of anthrax suitability for India and has found that this suitability is strongly associated with the wildlife-livestock interface as represented by the presence of livestock across the spectrum of the ecological niche of a key sentinel wildlife species, *E. maximus*. While this study cannot claim whether this association is due to the specific ecology of elephants in ecotones, or whether it is simply due to their capacity to function as important anthrax sentinels, we can claim that intervening efforts to separate wildlife and livestock at potential points of contact in the landscape may be an important step toward preventing the cycling of anthrax between wild and domestic animals. Moreover, while not as impactful as the wildlife-livestock interface, the concurrent influence of historical forest loss lends further support to the potential impact of anthropogenic ecotones in the ongoing transmission of anthrax in India. These findings highlight the potential benefits of a One Health approach to anthrax prevention and control, incorporating the expertise and spheres of influence of state veterinary, forest management, and human health services.

## Authors’ contributions

MW carried out methodological conceptualization, modelling, and writing the manuscript and editing drafts; SM carried methodological conceptualization, writing the manuscript and editing drafts; SH carried out experiential and methodological conceptualization, modelling, and writing the manuscript and editing drafts.

## Acknowledgments

None

## Supporting Information Captions

Figure S1. The distribution of the environmental layers used in the modeling of the anthrax niche. Distribution of anthrax outbreaks documented by the Pro-MED mail surveillance mechanism between 2000 and 2018 is also shown (black dots). All maps are displayed only for the purposes of depicting the distribution of disease occurrence and risk, and do not reflect the authors’ assertion of territory or borders of any sovereign country including India. All maps created in R (v. 3.3.1).

Figure S2. The distribution of predicted anthrax suitability from each model presented in S2 Table 1. All maps are displayed only for the purposes of depicting the distribution of disease occurrence and risk, and do not reflect the authors’ assertion of territory or borders of any sovereign country including India. All maps created in R (v. 3.3.1).

Figure S3. The Maxent model with the jackknife variable selection procedure comparing each covariate’s lone contribution to the training gain (blue) to its effect on training gain when the covariate is withheld from the model (red).

Figure S4. Predicted anthrax suitability with elephant outbreaks removed from the training data. All maps are displayed only for the purposes of depicting the distribution of disease occurrence and risk, and do not reflect the authors’ assertion of territory or borders of any sovereign country including India. All maps created in R (v. 3.3.1).

Table S1. Maxent model comparisons using the area under the receiver operating characteristic curves (AUC), the Akaike information criteria (AIC), and the covariate rankings. The AUC for each model is based on testing against an independent sample of laboratory-confirmed anthrax outbreaks, while the AIC is derived from a Poisson point process. Rankings are based on the permutation importance of each covariate and its contribution, reported as a percentage, to the loss function during the fitting of the Maxent model.

